# An iTSC-derived placental model of SARS-CoV-2 infection reveals ACE2-dependent susceptibility in syncytiotrophoblasts

**DOI:** 10.1101/2021.10.27.465224

**Authors:** J Chen, JA Neil, JP Tan, R Rudraraju, M Mohenska, YBY Sun, G Sun, Y Zhou, Y Li, D Drew, P Pymm, WH Tham, FJ Rossello, G Nie, X Liu, K Subbarao, JM Polo

**Author notes:** These authors contributed equally (J Chen, JA Neil, JP Tan, R Rudraraju).

## Abstract

Severe acute respiratory syndrome coronavirus 2 (SARS-CoV-2) infection causing coronavirus disease 2019 (COVID-19) has caused a global health crisis. The primary site of infection is in the respiratory tract but the virus has been associated with a variety of complications involving the gastrointestinal and cardiovascular systems. Since the virus affects a variety of tissue types, there has been interest in understanding SARS-CoV-2 infection in early development and the placenta. ACE2 and TMPRSS2, two genes that are critical for SARS-CoV-2 virus entry are expressed in placenta-specific cell types including extravillous trophoblasts (EVTs) and especially, syncytiotrophoblasts (STs). The potential of SARS-CoV-2 to infect these placental cells and its effect on placental development and function is still unclear. Furthermore, it is crucial to understand the possible mechanism of vertical transmission of SARS-CoV-2 through the placenta. Here, we developed an *in vitro* model of SARS-CoV-2 infection of placental cell types using induced trophoblast stem cells (iTSCs). This model allowed us to show that STs but not EVTs are infected. Importantly, infected STs lack the expression of key differentiation genes, lack typically observed differentiated morphology and produce significantly lower human chorionic gonadotropin (HCG) compared to non-infected controls. We also show that an anti-ACE2 antibody prevents SARS-CoV-2 infection and restores normal ST differentiation and function. We highlight the establishment of a platform to study SARS-CoV-2 infection in early placental cell types, which will facilitate investigation of antiviral therapy to protect the placenta during early pregnancy and development.

## Introduction

In 2020, a novel coronavirus called severe acute respiratory syndrome coronavirus (SARS-CoV-2) emerged and spread worldwide, infecting millions of people and causing Coronavirus Infectious Disease 2019 (COVID-19). COVID-19 was declared a public health emergency by the World Health Organization in February 2020 (Zhu et al., 2020). The infection is asymptomatic or mild in the majority of cases, however, patients can develop severe illness associated with acute respiratory distress syndrome (ARDS) and organ failure (Aguiar et al., 2020, Brosnahan et al., 2020, Long et al., 2020). This disease can lead to death in 26% of critically ill patients (Grasselli et al., 2020) to as high as 61.5% in another report (Yang et al., 2020). This has led to an imperative to understand the extent of cells that can be infected, the mechanism of infection and develop adequate treatments.

The cellular receptor for SARS-CoV-2 is angiotensin-converting enzyme 2 (ACE2) that is present in many cell types such as lung alveolar epithelial cells, enterocytes, venous endothelial cells, and smooth muscle cells (Bhalla et al., 2020, Hamming et al., 2004, Jing et al., 2020). Along with ACE2, transmembrane serine protease 2 (TMPRSS2) has been identified as an entry factor for SARS-CoV-2 infection. Cells that express both receptors are susceptible to infection (Hoffmann et al., 2020). Several reports have utilised *in vitro* derived systems to understand SARS-CoV-2 infection and screen for antiviral drugs or potential treatments. Conventionally, Vero cells, which have been derived from kidney epithelial cells of the African green monkey, have been used to study several viral infections *in vitro* (Govorkova et al., 1996, Ksiazek et al., 2003), and more recently one of the *in vitro* cell models used to replicate and isolate SARS-CoV-2 (Mantlo et al., 2020, Takayama, 2020, Zhou et al., 2020). However, these cells fail to recapitulate many aspects of infection in human cells. Therefore, in order to facilitate a more accurate model relevant to human infectivity, pluripotent stem cells (PSCs)-derived *in vitro* models of cardiac, kidney, lung and intestinal cells have been used as platforms to study SARS-CoV-2 (Bailey et al., 2021, Huang et al., 2020, Lamers et al., 2020, Monteil et al., 2020, Zhao et al., 2020). These models are especially valuable as they not only better represent the nuances of different human cell types but are scalable and tractable.

Despite advances in understanding SARS-CoV-2 infection in several somatic cell types, little is known about the virus and its impact on early development of the embryo or placenta. There have been clinical reports on the effects of SARS-CoV-2 in the early and late pregnancy that show increased risk of complications in infected patients (Garrido-Pontnou et al., 2021, Jang et al., 2021). In the placenta, SARS-CoV-2 infection was observed within the villi or intervillous space (Best Rocha et al., 2020, Hecht et al., 2020, Morotti et al., 2021, Patane et al., 2020). These reports observed that villous syncytiotrophoblasts were the primary target of infection based on histopathological evidence (Hecht et al., 2020, Morotti et al., 2021, Patane et al., 2020). In spite of the clinical evidence, the effect of SARS-CoV-2 infection on placental health and fetal development is still unclear. An *in vitro*-derived model is key to understanding the effects of SARS-CoV-2 infection in the placenta.

The derivation of trophoblast stem cells (TSCs) *in vitro* from first trimester placenta or blastocysts capable of differentiating into both main placental cell types: extravillous cytotrophoblasts (EVTs) and syncytiotrophoblasts (STs), facilitates the study of placenta biology and pathology *in vitro* (Okae et al., 2018). Furthermore, we and others recently reported that fibroblasts can be reprogrammed into induced trophoblast stem cells (iTSCs) which are molecularly and functionally similar to TSCs (Castel et al., 2020, Liu et al., 2020). Therefore, TSCs and iTSCs constitute a scalable and tractable model to study *in vitro* placental biology. Here we utilised iTSCs to generate an *in vitro* model of placenta infection by SARS-CoV-2. We found that STs were the only cell type that was productively infected, leading to phenotypic, transcriptomic and metabolic changes, with functional consequences. Importantly, we showed that infection and concomitant cellular changes in STs could be prevented using ACE2 antibodies.

## Results

### STs are productively infected with SARS-CoV-2

We first verified the expression of ACE2 in the 1st trimester placenta by immunohistochemistry. Trophoblast cells within the placental villi, especially STs lining the villous surface (marked by HCG staining), were strongly positive for ACE2 (Fig 1A). In the maternal decidua, multiple cells were also faintly stained for ACE2, including EVTs which were identified by HLA-G positivity on serial sections (Fig 1B). This is consistent with previous reports that STs and EVTs express ACE2 (Pringle et al., 2011, Valdes et al., 2006). We have previously reported the *in vitro* generation of induced trophoblast stem cells (iTSCs) from fibroblasts, which could be used to model placental cell differentiation into EVTs and STs (Liu et al., 2020). These EVTs and STs expressed typical cell-specific markers HLA-G and HCG respectively (Fig 1C). We examined the expression of ACE2 in iTSCs before and after differentiation into EVTs and STs in two independent donor cell lines (32F and 55F) (Fig 1C, Fig S1A-C), and detected ACE2 mRNA and protein in EVTs and STs but not in iTSCs (Fig 1C-D). EVTs and STs also expressed higher levels of TMPRSS2 relative to iTSCs (Fig 1D). The expression of these 2 important factors, particularly ACE2, in EVTs and STs suggests a potential susceptibility to SARS-CoV-2 infection.

**Figure 1.**
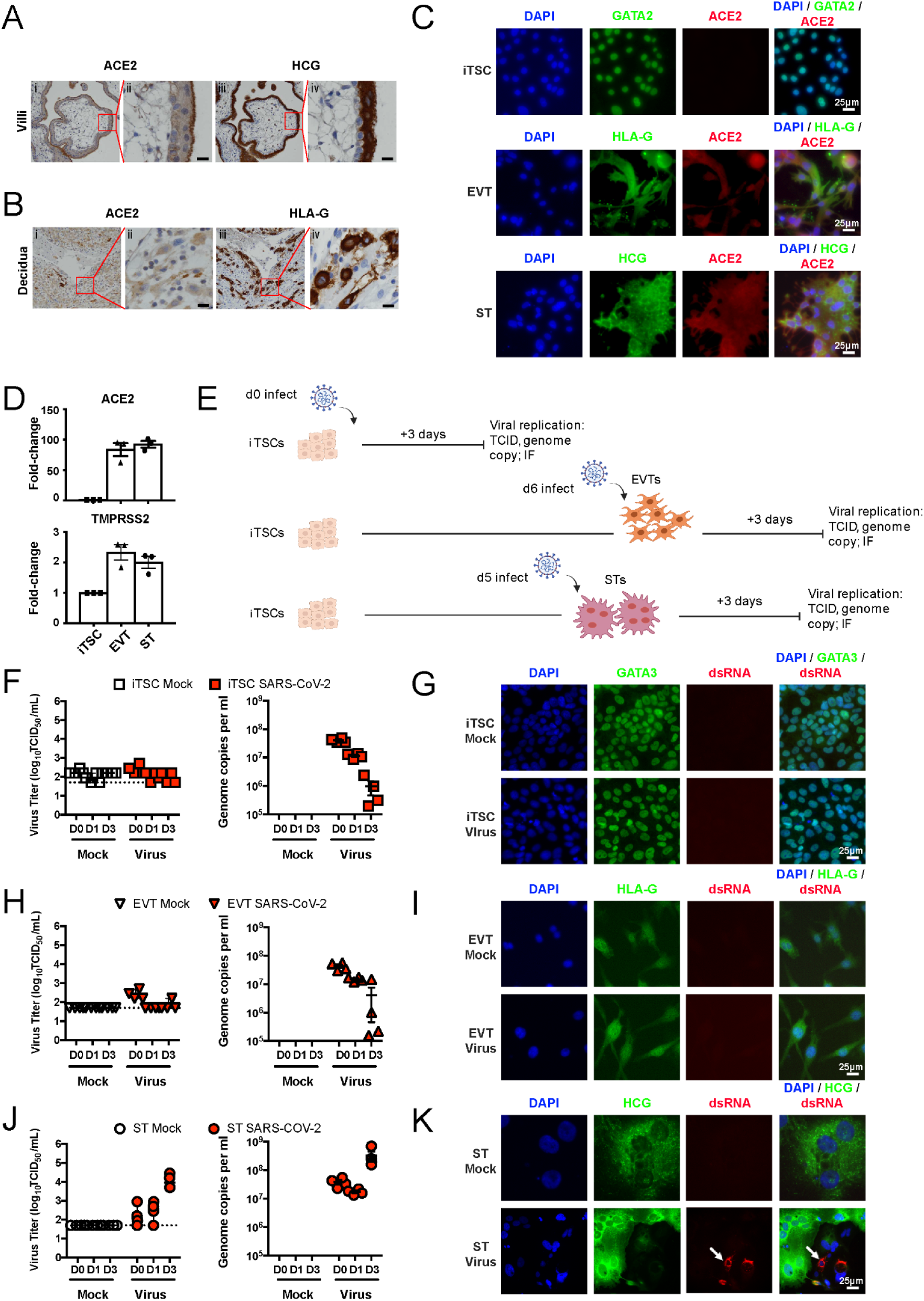
STs are susceptible to SARS-CoV-2 infection. A) Immunohistochemistry images of 1^st^ trimester placental villi for ACE2 and HCG. Scale bar = 200µm. B) Immunohistochemistry images of maternal decidua for ACE2 and HLA-G. Scale bar = 200µm. C) Immunofluorescence images of ACE2(Red) along with GATA3 (TSCs; Green), HLA-G (EVTs; Green) or HCG (STs; Green). D) qPCR analysis of ACE2 and TMPRSS2 expression in TSCs, EVTs and STs (fold-change relative to TSCs). E) Schematic of differentiation and infection with SARS-CoV-2. Virus titre expressed in log_10_TCID_50_/ml, genome copy analysis, and Immunofluorescence images of dsRNA in 55F TSCs (E, F), EVTs (G, H) and STs (I, J). Cells counterstained with DAPI. n=3 (A); n=3 (D); n=4 technical replicates of a representative experiment of 32F (n=2) and 55F (n=2) lines (F, H, J).

In order to test if these cells were susceptible to infection, we infected iTSCs, as well as EVTs and STs towards their terminal differentiation at day 6 and day 5 respectively (Fig 1E). In iTSCs and EVTs, no significant increase in viral titres was observed in the supernatant over time and intracellular dsRNA was not detected at day 3 post infection, which would have been evidence of active viral replication within the cells (Valdespino-Vazquez et al., 2021) (Fig 1F-I). In contrast, in STs viral titres increased in the supernatant 3 days after infection (Fig 1J); infection was confirmed by identification of dsRNA within ST cells (Fig 1K), which was similar to a report on SARS-CoV-2 infection of placental tissue (Hosier et al., 2020).

### SARS-CoV-2 affects differentiation potential and metabolic activity

Since STs are the only cell type that was infected by SARS-CoV-2 in our model, we focused on this cell type for further study and dissected dynamics of SARS-CoV-2 infection in-depth. Placental development is critically dependent on trophoblast cell differentiation into EVTs and STs (Arnholdt et al., 1991); therefore, we wanted to understand how early during differentiation STs could be infected. We first analysed the expression of ACE2 during the differentiation of iTSCs into STs and found that cells begin to express ACE2 as early as day 2 of differentiation (Fig 2A). We verified this increase of expression by immunofluorescent staining for ACE2 in both cell lines (Fig S1A and B). To test whether the expression of ACE2 was sufficient to confer susceptibility to infection by SARS-CoV-2, we infected STs at days 2, 3, and 4 of differentiation (Fig 2B) and found that cells could be infected as early as day 2 of differentiation (Fig 2C). Positive/productive infection was identified by the presence of dsRNA within the cells and presence of infectious virus in the cell supernatant in both cell lines (Fig S2C). In addition, we observed that the differentiation potential of STs was affected as infected (dsRNA+) cells appeared morphologically more immature (please see methods) than non-infected (dsRNA-) cells and had lower signal of HCG despite being multinucleated, suggesting that SARS-CoV-2 infection severely affects cellular differentiation (Fig S2D). We further assessed levels of HCG in cells and cellular morphology, to quantify proportions of differentiated and undifferentiated cells. We then categorized differentiated and undifferentiated cells as either dsRNA+ or dsRNA- to determine which cells were infected. We found that the ST cells that were dsRNA+ within a virus-infected culture (cells exposed to the virus and were indeed infected) were significantly less differentiated than dsRNA-cells (cells exposed to the virus but were not infected) within the same culture (Fig 2D). Importantly, dsRNA- cells within the virus-infected cultures had a similar differentiation potential as mock-infected cells. Taken together with the result seen in dsRNA+ cells, the data suggest that SARS-CoV-2 infection hinders the differentiation of STs.

**Figure 2.**
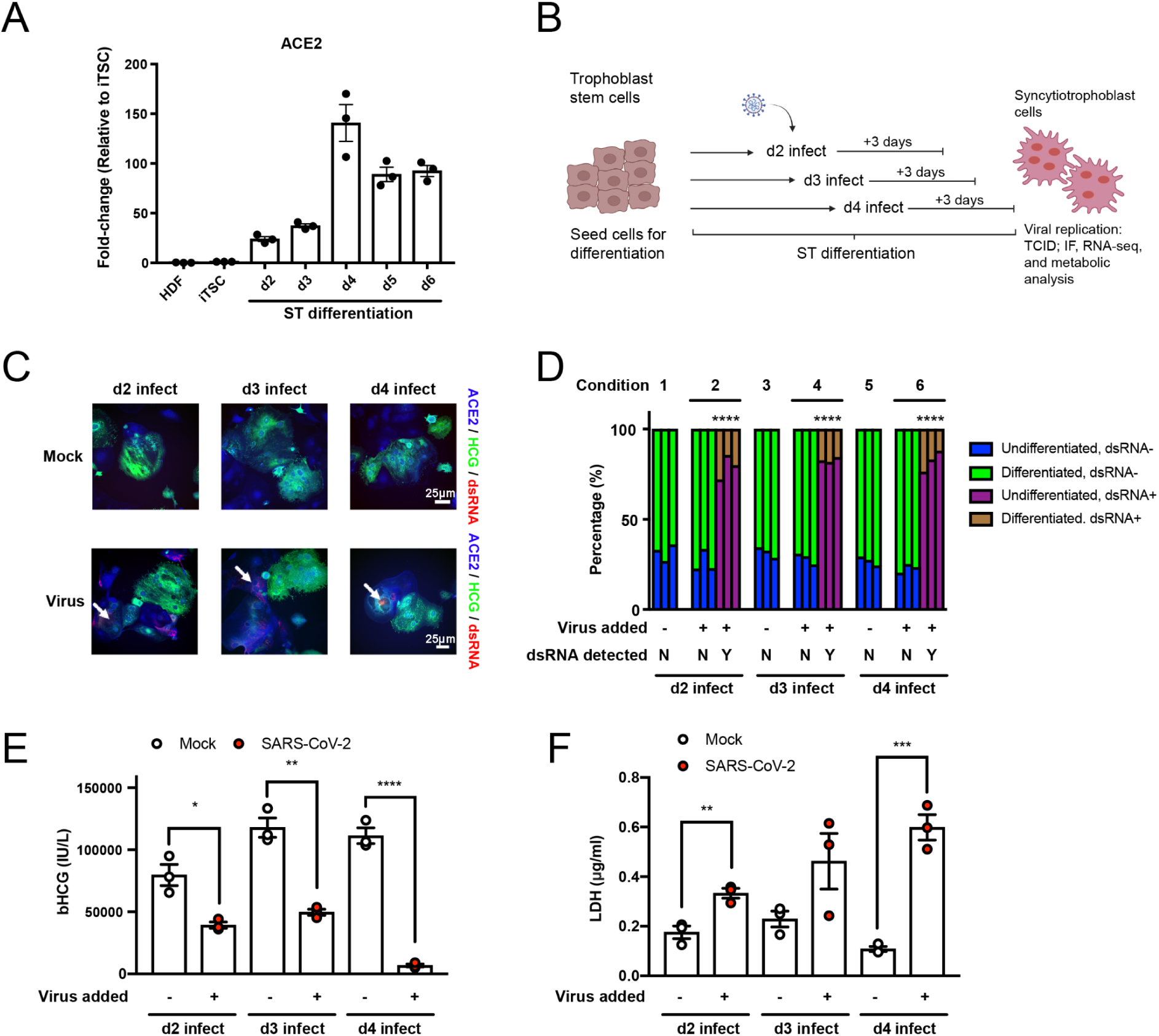
SARS-CoV-2 can infect STs during differentiation and affects differentiation potential and metabolic activity. A) qPCR analysis of ACE2 expression of ST differentiation (d2-6) (fold-change relative to TSCs). B) Schematic of infection of STs during differentiation. C) Immunofluorescence images of dsRNA (Red) and HCG (Green) in 55F STs of mock and virus infected conditions (d2, 3, and 4). D) Quantification of cellular differentiation of dsRNA+/dsRNA- cells in infected or mock conditions. E) bHCG assay of 55F STs during differentiation. F) LDH assay of 55F STs during differentiation. Cells counterstained with DAPI. n=3 (A and D); n=3 technical replicates of a representative experiment (E and F). *p<0.05, **p<0.01, ***p<0.005, ****p<0.001

Since STs produce human chorionic gonadotropin (HCG), the hormone that is vital for maintaining pregnancy, we analysed the metabolic activity in the cultures during differentiation (Nwabuobi et al., 2017). We observed that HCG levels were significantly lower in infected cells throughout differentiation (day 2, 3 and 4) compared to mock controls (Fig 2E). We also analysed cell death using a lactate dehydrogenase (LDH) assay and found that infected cells released higher levels of LDH in the supernatant compared to mock controls, indicating significantly more cell death in the SARS-CoV-2infected cultures (Fig 2F). Taken together, SARS-CoV-2 infection of STs leads to an impairment of differentiation potential, lack of HCG production and increased cell death during differentiation.

### Transcriptome-wide profiles of infected STs show an increase in viral responses

To understand the effects of infection on the cells in greater depth, we analyzed the transcriptome of ST cells infected at day 3 of differentiation and 3 days post-infection to reach the theoretical fully differentiation time course (day 6), compared to mock-infected cells. Correspondence analysis (CoA) indicated that SARS-CoV-2-infected cells were transcriptionally divergent from mock-infected cells (Fig 3A). Differential gene expression (DGE) analysis identified 155 genes upregulated and 140 genes downregulated in infected cells (Table S1). Importantly, we identified that among the differentially expressed genes (DEGs), ST-specific genes such as CGA and PSG3 were significantly down regulated in SARS-CoV-2 infected cells compared to mock-infected cells, suggesting cultures were not differentiating into STs as efficiently as were mock-infected controls. Upregulated genes were related to interferon signaling (IFNL1, IFNB1, IFIH1) (Fig 3B). Other genes associated with TNFα signaling via NFKB such as MAPK4, STAT1, RELB and NFKBIA were also highly upregulated in infected cells compared to mock-infected cells, indicating that there was a strong innate response to viral infection (Mantlo et al., 2020). Furthermore, gene ontology analysis showed an enrichment of viral response, along with antiviral mechanism (IFN-stimulation) and response to type I interferon in the significantly upregulated genes in infected cells. In contrast, genes downregulated upon infection were enriched in cellular and metabolic processes such as Asparagine N-linked glycosylation, and response to endoplasmic reticulum stress (Fig 3C). Although not strongly enriched, we did identify 14 significantly up-regualted DEGs involved in positive regulation of cell death such as, EGR1, FOS, SNCA and PHLDA1. Moreover, expression levels of TSC, EVT and ST identity-specific genes, showed that while mock-infected cells have robust expression of ST-related differentiation genes, SARS-CoV-2 infected cultures failed to upregulate ST associated genes, and infected cultures also had higher levels of expression of TSC-related genes; suggesting that cells were less differentiated, consistent with our observations that infected cultures have impaired differentiation potential (Fig 3D). Taken together, we show that SARS-CoV-2 infection of ST cultures elicits an NFKB-mediated inflammatory response and has a negative impact on the differentiation pathway of cells.

**Figure 3.**
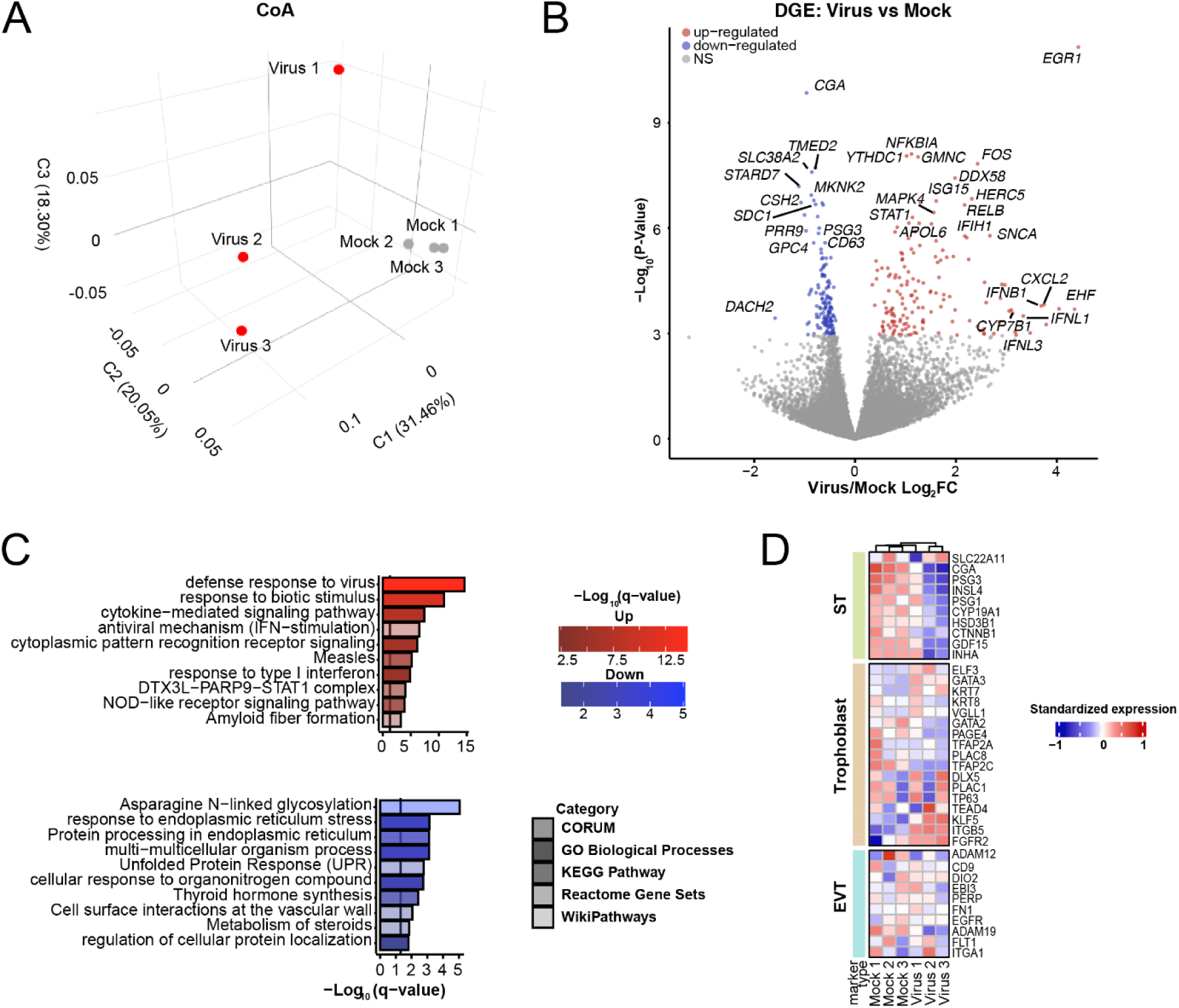
SARS-CoV-2 impacts the transcriptomic profile in STs and upregulates innate inflammatory response to infection. A) Transcriptome-wide CoA of virus and mock conditions of d3 infected STs. B) Volcano plot depicting DGE analysis between virus and mock conditions. C) Functional enrichment analysis of significantly upregulated or downregulated gene sets (FDR <0.05). D) Hierarchically clustered heatmap of cell identity genes expressed in STs, under virus and mock conditions.

### Inhibition of ACE2 prevents viral infection of STs

Finally, as the high expression of ACE2 seemed to correlate well with the susceptibility of STs to SARS-CoV-2, we investigated whether ACE2 could be targeted to inhibit SARS-CoV-2 entry into STs using an anti-ACE2 antibody. We generated and characterised antibodies against recombinant human ACE2 from a phage library (Fig S3A). We then validated the binding affinity of these clones and utilized them in our antibody blocking experiments (Figure S3B-C). Among the clones validated, we selected WCSL141 and WCSL148 for the blocking experiments. We show that SARS-CoV-2 infection of STs was blocked by the addition of two anti-ACE2 antibodies (Virus^αACE2^) (Fig 4A). We did not detect infectious virus or viral genome copies in culture supernatants while both were detected in virus-treated control antibody conditions (Virus^Ctrl^) demonstrating that ACE2 antibody inhibits virus entry (Fig 4B). We further analysed the transcriptomes of cells from these separate conditions. Hierarchical clustering of samples showed that cultures blocked with anti-ACE2 antibody (Virus^aACE2^) clustered closely with other mock-infected samples and were separate from infected cells treated with a control antibody (Virus^Ctrl^) (Fig 4C). CoA also showed a divergent transcriptome of cells from the Virus^Ctrl^ condition from the rest (Fig S4A). In addition, we also verified SARS-CoV-2 expression in the samples. As expected, we observed high expression in Virus^Ctrl^, and neither in uninfected cells, nor, surprisingly, in Virus^aACE2^. (Fig S4B, S4C). We further verified our results by performing unsupervised k-means clustering of all genes across the samples, and identified unique clusters of genes that were upregulated and downregulated in Virus^Ctrl^ in contrast to the other samples (Fig 4D, Table S1). Interestingly, we did not identify a specific signature in the treated and infected cells (Virus^aACE2^). We then performed functional enrichment analysis on clusters 1 and 2, which contained strongly upregulated or downregulated in infected samples (Virus^Ctrl^). As before, we found an upregulation of host-defense response to virus and IFN signalling pathways, as well as downregulation of cellular and metabolic processes, such as transport of small molecules and nucleobase-small molecule metabolic process (Fig S4D). We explored placental identity genes and found that Virus^aACE2^ cultures exhibited an increase in expression of ST identity genes, similar to mock-infected controls (Fig 4E). As the expression levels of differentiation-specific genes was similar to mock controls , we showed that anti-ACE2 antibody blocking rescued the differentiation potential of cells compared to SARS-CoV-2 infected cells, resulting in a similar number of cells with morphological features akin to mock-infected controls (Fig 4F). We went on to assess whether the restoration of differentiation potential would also lead to a restoration of metabolic processes in STs. We observed that Virus^aACE2^ cultures showed comparable production of HCG to mock-infected controls (Fig 4G) and lower levels of LDH compared to Virus^Ctrl^ cells(Fig 4H). Taken together, we demonstrated that SARS-CoV-2 infects ST cells and affects their differentiation and metabolic activity. Cells could differentiate and were metabolically active when infection was prevented by blocking ACE2.

**Figure 4.**
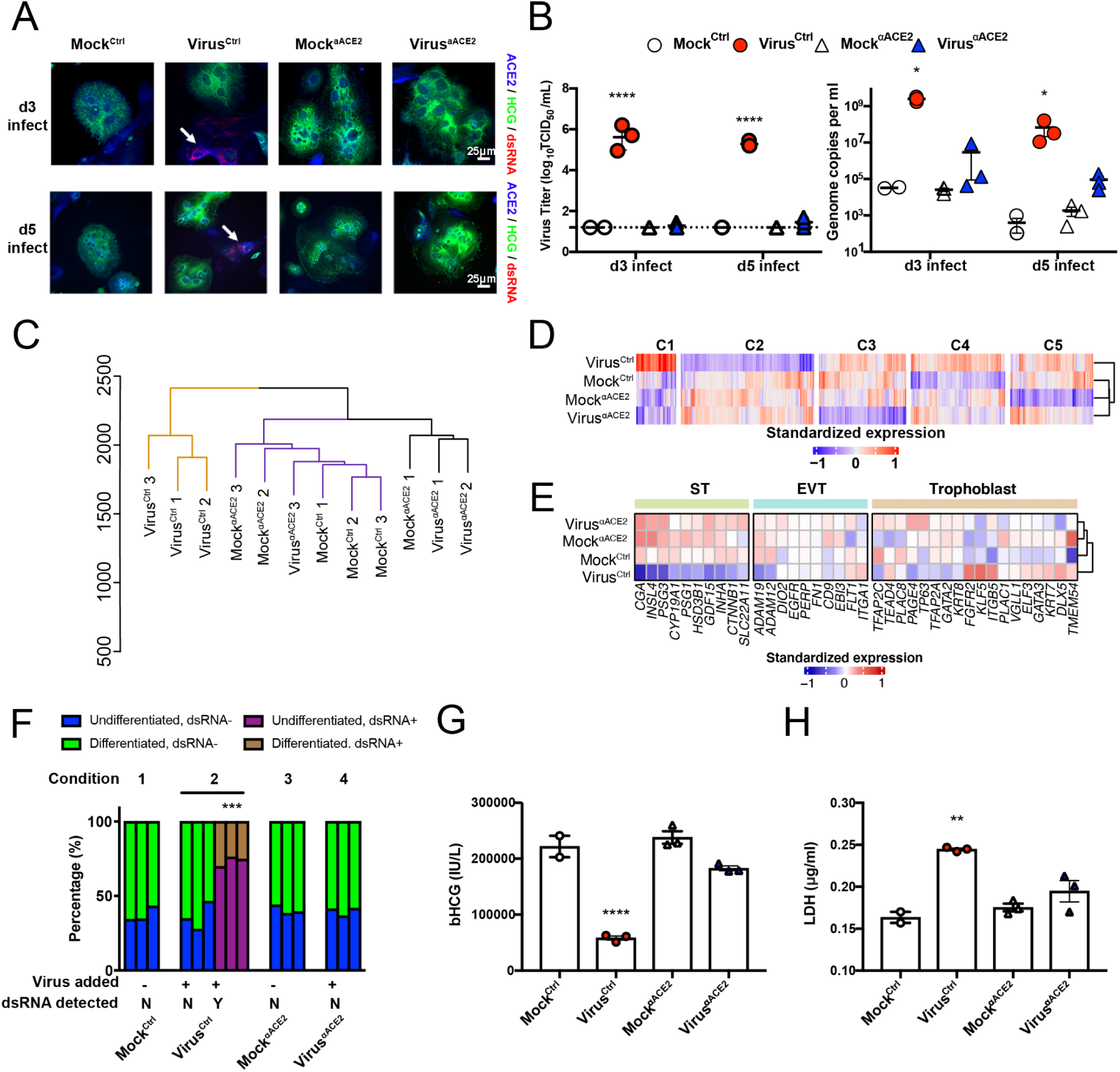
Inhibition of ACE2 via anti-ACE2 antibody restores normal differentiation and function of STs. A) Immunofluorescence images of dsRNA (Red) and HCG (Green) in STs of anti-ACE2 antibody and/or virus treated conditions at d3 and d5. Cells counterstained with DAPI. B) Virus titre expressed in log_10_TCID_50_/ml and genome copy analysis of antibody and/or virus treated conditions at day 3 and day 5. C) Transcriptome-wide hierarchical clustering analysis of d3 infected/mock/treated STs. D) Transcriptome-wide k-means clustering analysis of d3 infected/mock/treated STs; cluster 6 contained genes with no distinct pattern between the samples and is not shown E) Hierarchically clustered heatmap of cell identity genes expressed in STs, under virus, mock and treated conditions) .F) Quantification of cellular differentiation of dsRNA+/dsRNA- STs in infected/mock conditions. G) HCG assay in STs antibody/virus conditions. H) LDH assay of STs in antibody/virus conditions. n=3 experimental replicates (B-H; except n=2 in d3 Mock^Ctrl^ in B). n=3 technical replicates of a representative experiment (B, G, H; except n=2 in d3 Mock^Ctrl^ in B). *p<0.05, **p<0.01, ***p<0.005, ****p<0.001

## Discussion

The use of *in vitro* models of different cell types to study SARS-CoV-2 infection allows us to understand inherent mechanisms. It also enables us to perform and test specific drug treatments for affected cell types (Han et al., 2020, Huang et al., 2020, Pei et al., 2020). In our *in vitro* model, SARS-CoV-2 is able to infect STs but not iTSCs or EVTs. While disparities between *in vitro* culture systems and clinical specimens have been observed in the lung with SARS-CoV-2 infected alveolar type 1 (AT1) cells observed in organoids but not isolated lung tissue (Pei et al., 2020, Bradley et al., 2020), our data are consistent with histopathological studies using clinical specimens. These indicate that STs in the intervillous space are the typical cells that harbor SARS-CoV-2 in infected placentas (Garrido-Pontnou et al., 2021, Hecht et al., 2020, Jang et al., 2021, Morotti et al., 2021, Patane et al., 2020). Although EVTs were not infected by SARS-CoV-2 in our model, others have reported that these cells are susceptible to viruses like adenovirus(Koi et al., 2001) . The reason for lack of SARS-CoV-2 infection in the in vitro generated EVTs despite evidence of ACE2 and TMPRSS2 expression is unclear and will require further investigations.

SARS-CoV-2 was able to infect STs during differentiation, which led to cell death and impaired differentiation. Since iTSC derived STs are a model of early placentation, these results support clinical evidence that the virus could affect the placenta in early development (Valdespino-Vazquez et al., 2021). Furthermore, the sensitivity and tractability of our *in vitro* ST model enabled the assessment of functional impairments resulting in reduced levels of HCG production. Reduced HCG production may be associated with complications in pregnancy, including early miscarriage (Jing et al., 2020). We were also able to determine the phenotypic effects of the virus on STs, which corresponded transcriptionally to the lack of expression of genes typical of ST differentiation and identity. Consistent with other reports of SARS-CoV-2 infection in a variety of cell types, we identified an upregulation of viral response and innate immunity genes. Importantly, SARS-CoV-2 infection elicited an IFN response in our model, similar to responses in other cell types such as lung and cardiac cells (Huang et al., 2020, Li et al., 2021, Mulay et al., 2021). Downregulation of genes involved in cellular processes and function were also observed (Daniloski et al., 2021, Suryawanshi et al., 2021). Although, as demonstrated by our results, this infection model can be of great utility, there are limitations. For example, the decidua contains 10-20% of macrophages but we cannot model their effects in this system (Kreis et al., 2020, Manaster and Mandelboim, 2010). In the future, complex models that include immune cells could be used to enhance current models, as has been done with brain slice cultures and microglia from iPSC derived models (Grubman et al., 2021). Recent studies have reported that host cell factors such as ACE2, TMPRSS2 or cathepsins are vital for SARS-CoV-2 entry and could be utilized as potential therapeutic targets against infection (Dong et al., 2020, Hoffmann et al., 2020). We also reported that blocking viral infection through ACE2 blockade restores the functional phenotype in STs, similar to the rescue of function in lung and cardiac cells through the inhibition of ACE2 or TMPRSS2 activity (Huang et al., 2020, Hoffmann et al., 2021, Pei et al., 2020, Bojkova et al., 2020, Perez-Bermejo et al., 2021). More importantly, this *in vitro* derived placental model allowed us to generate a quick and effective system. We envision that our model will facilitate a deeper understanding of COVID-19 pathogenesis and will provide a platform for drug discovery and potential treatments.

## Author Contributions

J.M.P and K.S. conceptualised and supervised the study. J.C. and J.P.T. performed all the cellular work with the support of YBY.S., G.S and Z.Y.. J.A.N. and R.R performed all the viral work. J.C., J.P.T., J.A.N and R.R performed all the molecular and microscopic analysis in the infected cells. M.M performed the bioinformatic analysis with the supervision of F.J.R and J.M.P.. Y.L. performed the placenta staining under the supervision of G.N.. D.D., P.P., and W.H.T generated, characterized and provided the anti-ACE2 antibodies. J.M.P, K.S., J.C., J.P.T., J.A.N and R.R wrote the manuscript with contributions of all the authors.

## Acknowledgments

The authors thank the ACRF Centre for Cancer Genomic Medicine at the MHTP Medical Genomics Facility for assistance with next generation library preparation and Illumina sequencing. J.M.P. and K.S. were supported by a MRFF grant (MRF9200007) and a DHHS Victorian Government Grant. J.M.P was supported by an ARC Future Fellowship as a NHMRC Ideas Grant (APP2004774). K.S. is supported by an NHMRC Investigator grant (APP1177174). The Melbourne WHO Collaborating Centre for Reference and Research on Influenza is supported by the Australian Government Department of Health. W-HT is a Howard Hughes Medical Institute–Wellcome Trust International Research Scholar (208693/Z/17/Z) and anti-ACE2 antibody generation was supported by the Victorian Government and the Medical Research Future Fund (MRFF) GNT2002073.

## Materials and Methods

### Placental tissue preparation and Immunohistochemistry

Ethics approval for the use of first trimester human placental tissues for research was obtained from the Human Ethics Committee at Monash Health (RES-19-0000-399A, Melbourne, Australia), all patients provided informed written consent. Tissues were collected after elective pregnancy termination, fixed in buffered formalin and embedded in paraffin.

Tissues were sectioned (5um) onto SuperFrost® Plus slides (J3800AMNT, ThermoFisher Scientific, MA, USA), dried overnight at 37°C, deparaffinized in histolene, then rehydrated in graded solutions of ethanol to Milli-Q water. Following antigen retrieval by microwaving for 10 min in 0.01M citrate buffer (pH 6.0), endogenous peroxidase was quenched with 3% H_2_O_2_, and tissues were next incubated with a blocking buffer containing high salt TBS (0.3 M NaCl in 50 mM Tris, pH 7.6), 0.1% Tween 20 (P2287, Sigma-Aldrich, USA), and 15% goat or horse serum for 20 min at room temperature (RT). Sections were then incubated with primary antibodies (ACE2, ab15438, Abcam, UK; HCG beta, ab9582, Abcam, UK; HLA-G, 557577, BD Pharmingen) for 1 h at 37°C; rabbit or mouse IgG (X0936 and X0931 respectively, Dako, USA) was used in negative controls. Sections were next incubated first with a biotinylated goat anti-rabbit IgG or horse anti-mouse IgG (BA-1000 and BA-2000 respectively, Vector Laboratories, USA), then with an avidin-biotin-complex conjugated to horseradish peroxidase (PK-6100, Vector Laboratories), each for 30 min at RT. All antibodies were incubated in blocking buffer containing TBS, 0.1% Tween 20 and 10% FCS. Color was developed with peroxidase substrate 3,3’-diaminobenzidine (DAB) (K3466; Dako). Sections were counterstained with Harris hematoxylin (HHS16, Sigma-Aldrich), mounted in DPX new mounting medium (100579, Sigma-Aldrich), and imaged using an Olympus microscope fitted with a Fujix HC-2000 high resolution digital camera (Fujix, Tokyo, Japan).

### Cell culture and differentiation

Induced trophoblast stem cells (iTSCs) were maintained as previously (Liu et al., 2020, Okae et al., 2018). Briefly, iTSCs were cultured in TSC medium consisted of DMEM/F-12, GlutaMAX (ThermoFisher) supplemented with 0.3% BSA (Sigma), 0.2% FBS (ThermoFisher), 1% ITS-X supplement (ThermoFisher), 0.1 mM 2-mercaptoethanol (ThermoFisher), 0.5% penicillin–streptomycin (ThermoFisher), 1.5 μg/ml l-ascorbic acid (Sigma), 5 μM Y27632 (ROCK inhibitor; Selleckchem), 2 μM CHIR99021 (Miltenyi Biotec), 0.5 μM A83-01 (Sigma), 1 μM SB431542, 50 ng/ml EGF (Peprotech) and 0.8 mM VPA (Sigma) onto 5 μg/ml collagen IV (Sigma-Aldrich)-coated plates. The iTSCs were routinely passaged every 4-5 days with medium replacement performed every other day.

Differentiation of iTSCs into STs and EVTs was performed and modified as previously described (Okae et al., 2018). For the differentiation of iTSCs into STs, iTSCs were seeded at a density of 3.75 × 10^4^ cells per well onto a 24-well plate pre-coated with 2.5 μg/ml collagen IV (Sigma) and cultured in 500 μl ST differentiation medium (DMEM/F-12, GlutaMAX (ThermoFisher) supplemented with 0.3% BSA (Sigma), 4% KSR (ThermoFisher), 1% ITS-X supplement (ThermoFisher), 0.1 mM 2-mercaptoethanol (ThermoFisher), 0.5% penicillin–streptomycin (ThermoFisher), 2.5 μM Y27632 (Selleckchem) and 2 μM forskolin (Selleckchem)). Medium was replaced on day 3 of differentiation, and cells were analysed on day 6. For the differentiation of iTSCs into EVTs, iTSCs were seeded at a density of 3.4 × 10^4^ cells per well onto a 24-well plate pre-coated with 1 μg/ml collagen IV (Sigma) and cultured in 500 μl EVT differentiation medium (DMEM/F-12, GlutaMAX (ThermoFisher) supplemented with 0.3% BSA (Sigma), 4% KSR (ThermoFisher), 1% ITS-X supplement (ThermoFisher), 0.1 mM 2-mercaptoethanol (ThermoFisher), 0.5% penicillin–streptomycin (ThermoFisher), 2.5 μM Y27632 (Selleckchem), 100 ng/ml hNRG1 (Cell Signaling), 7.5 μM A83-01 (Sigma) and 2% Matrigel (Corning). On day 3 of differentiation, the medium was replaced with EVT differentiation medium without hNRG1, and Matrigel (Corning) was added to a final concentration of 0.5%. On day 6 of differentiation, EVT differentiation medium was replaced without hNRG1 (Cell Signaling) and KSR (ThermoFisher), and Matrigel (Corning) was added to 0.5% final concentration. The cells were cultured for an additional 2 days before analyses were performed.

### SARS-CoV-2 infection

Placental cells (TSCs, terminally differentiated EVTs/STs and differentiating STs) in 24 well plates were infected in duplicate or triplicate with 10^4^ tissue-culture infectious dose 50 (TCID_50_) of SARS-CoV-2 (Australia/VIC01/2020) for 1h. Virus was removed and cells cultured in cell type-specific medium for 3 days. Supernatants were collected and medium replaced daily. Median TCID_50_ in supernatants were determined by 10-fold serial dilution in Vero cells and calculated using the Reed and Muench method. RNA was extracted from supernatants using the QIAamp Viral RNA mini kit (Qiagen) and E-gene expression assessed using the SensiFAST Probe No-Rox One Step Kit (Bioline) and the following primers/probes: Fwd: 5’-ACAGGTACGTTAATAGTTAATAGCGT’-3, Rev: ATATTGCAGCAGTACGCACACA and Probe: FAM-ACACTAGCCATCCTTACTGCGCTTCG-BBQ. Viral genomes were interpolated using a standard curve generated by a plasmid containing the E-gene. Each experiment was repeated independently at least twice.

### Phage Library Isolation of anti-ACE2 mAbs

Biopanning for anti-ACE2 human antibodies using the CSL human antibody phage library was performed as previously described (Panousis et al., 2016). Phages displaying human Fabs were enriched after three rounds of biopanning on biotinylated recombinant human ACE2 immobilised to streptavidin Dynabeads (Dynal M-280, Invitrogen, cat # 112.06D). After the third round of panning, individual clones were selected for further analyses by ELISA for the presence of human ACE2 binding phage. Positive clones were sequenced and annotated using the International ImMunoGeneTics database (IMGT) and aligned in Geneious Prime. Fabs from positive phage were reformatted into IgG1 expression plasmids and used for transient expression in Expi293 cells. Human IgG1 antibodies were purified using Protein-A affinity chromatography.

### Assessment of human antibody binding specificity by ELISA

96-well flat-bottomed MaxiSorp plates were coated with 50 μl of 125 nM recombinant human or mouse ACE2 protein in PBS at room temperature for one hour. All washes were done three times using PBS and 0.1% Tween (DPBS-T) and all incubations were performed for one hour at room temperature. Coated plates were washed and blocked by incubation with 4% skim milk solution. Plates were washed and then incubated with 50 μl of 125 nM of anti-ACE2 mAbs. The plates were washed and incubated with horseradish peroxidase (HRP)-conjugated Goat anti-Human IgG secondary antibody (1:5000). After a final wash, 50 μL of azino-bis-3-ethylbenthiazoline-6-sulfonic acid (ABTS liquid substrate; Sigma) was added and incubated in the dark at room temperature for 20 minutes and 50 μL of 1% SDS was used to stop the reaction. Absorbance was read at 405 nm and all samples were done in duplicate.

### Affinity measurements using bio-layer interferometry

Affinity determination measurements were performed on the Octet RED96e (FortéBio). Assays were performed at 25 °C in solid black 96-well plates agitated at 1000 rpm. Kinetic buffer was composed of PBS pH 7.4 supplemented with 0.1% (w/v) BSA and 0.05% (v/v) TWEEN-20. All assays were performed using anti-human IgG Fc capture sensor tips (AHC) sensors (FortéBio). A 60 s biosensor baseline step was applied before anti-ACE2 mAbs (5 mg/mL) were loaded onto AHC sensors. For affinity measurements against human ACE2, antibodies were loaded by submerging sensor tips for 200 s and then washed in kinetics buffer for 60 s. Association measurements were performed by dipping into a two-fold dilution series of human ACE2 from 6 - 200 nM for 180 s and dissociation was measured in kinetics buffer for 180 s. Sensor tips were regenerated using a cycle of 5 s in 10 mM glycine pH 1.5 and 5 s in kinetic buffer repeated five times. Baseline drift was corrected by subtracting the average shift of an antibody loaded sensor not incubated with protein and an unloaded sensor incubated with protein. Curve fitting analysis was performed with Octet Data Analysis 10.0 software using a global fit 1:1 model to determine K_D_ values and kinetic parameters. Curves that could not be fitted were excluded from the analyses. Mean kinetic constants and standard error of the mean reported are the result of three independent experiments.

### ACE2 Blockade

For ACE2 blockade, placental cells were treated with either 20 μg/ml of both WCSL141 and WCSL148 or 40 μg/ml of human IgG1 isotype control for 30 mins prior to SARS-CoV-2 infection as above. Following virus removal, cells were cultured in media containing 20 μg/ml of both ACE2-blocking antibodies or 40 μg/ml of isotype control until the end of the experiment. Infectious virus titres and genome copies were determined as above. Each experiment was repeated independently twice.

### bHCG and LDH detection

Supernatants collected on day 3 post infection were analysed for HCG and LDH levels. bHCG was measured using the Abnova HCG ELISA Kit (Cat no. KA4005) as per manufacturer’s instructions. All supernatants were diluted 1/2000 prior to analysis. LDH was measured using the Abcam LDH cytotoxicity kit II (Cat no. ab65393) as per manufacturer’s instructions in undiluted supernatants using a LDH standard curve.

### Immunofluorescence

Cultured cells were fixed in 4% PFA (Sigma Aldrich) in PBS for 10 min and then permeabilized in PBS containing 0.3% Triton X-100 (Sigma Aldrich). Cultures were then incubated with primary antibodies followed by secondary antibodies (see dilutions below). 4’,6-diamidino-2-phenylindole (DAPI) (1:1000) (Invitrogen, Thermo Fisher Scientific) was added to visualize cell nuclei. Images were taken with a DMi8 inverted microscope (Leica). Primary antibodies used in the study were: anti-HCG (ab9582, abcam, 1:200), anti-dsRNA (MABE1134, Merck, 1:200), anti-ACE2 (ab15348, abcam, 1:200), anti-GATA3 (MA1-028, Invitrogen, 1:100), and anti-HLA G (ab7759, abcam, 1:50). Secondary antibodies used in the study (all 1:400) were Alexa Fluor 488 goat anti-mouse IgG1 (A21121, Invitrogen, Thermo Fisher Scientific), Alexa Fluor 555 goat anti-Rabbit IgG (A21428, Invitrogen, Thermo Fisher Scientific), Alexa Fluor 555 goat anti-mouse IgG2a (A21137, Invitrogen, Thermo Fisher Scientific), Alexa Fluor 647 donkey anti-rabbit IgG (A31573, Invitrogen, Thermo Fisher Scientific).

### Image Analysis/Cell Quantification

Cell quantification was performed using the particle analysis option of the ImageJ software (http://rsb.info.nih.gov/ij/). Four fields of view taken at 10x magnification were scored first for DAPI-positive nuclei, followed by quantification of HCG and dsRNA positive cell bodies. All data were analyzed by one-way analysis of variance followed by Bonferroni’s post-hoc-test to calculate p values. Data were quantified from a total of three independent experiments.

### RNA extraction and qPCR

RNA was extracted from cells using RNeasy micro kit (74004, Qiagen) and QIAcube (Qiagen); or miRNeasy micro kit (217084, Qiagen) according to the manufacturer’s instructions. Reverse transcription was then performed using QuantiTect reverse transcription kit (Qiagen, catalogue no. 205311). Real-time PCR reactions were set up in triplicates using QuantiFast SYBR Green PCR Kit (Qiagen) and then carried out on the 7500 Real-Time PCR system (ThermoFisher) using LightCycler 480 software.

### Gene expression analyses

#### Pre-processing RNA-seq

Raw next generation RNA sequencing (RNA-seq) reads were obtained in FASTQ format, and prior to demultiplexing the forward read FASTQ was trimmed with trimmomatic to 18 nucleotides (nt) (the targeted read length as described above) with the following parameters: SE -phred33 CROP:18 MINLEN:18 (Bolger et al., 2014). FASTQ files were then demultiplexed with sabre (Joshi, 2011) with the parameters pe -c -u -m 1 -l 10 -n for the barcode indexes as stated above. Following this, demultiplexed sample reads were filter-trimmed with trimmomatic to the targeted read length of 101 nt, with the parameters SE -phred33 CROP:101 MINLEN:10 (Bolger et al., 2014). Sequencing reads were then mapped to a customised genome, composed of both GENCODE’s GRCh38.p13 and human SARS-CoV2 (RefSeq - NC_045512.2; see “Custom genome for mapping” below for further details), with STAR v2.5.2b (Dobin et al., 2013) and the parameters: --outSAMattributes All --alignIntronMax 1000000 --alignEndsType Local. Aligned BAM files were then sorted and indexed with sambamba (Tarasov et al., 2015) using default parameters; followed by deduplication by unique molecular identifiers (UMIs) using Je’s (v1.2) je markdupes function, with parameters: MM=0 REMOVE_DUPLICATES=true ASSUME_SORTED=true (Girardot et al., 2016). Read counts were then generated with Subread’s (v1.5.2) featureCount function (Liao et al., 2013), using default parameters.

### Gene expression analyses of the human genome

For each set of analyses (STs infected with virus, STs infected with virus and treated aACE2), genes mapped to the hSARS-CoV2 were first removed, and following this genes with low counts were filtered out. Specifically, genes with less than 5 raw read counts across all samples were removed, and genes with at least 1 count per million (CPM) in a minimum of 2 samples were kept. Prior to library size normalisation, normalisation factors were calculated with EdgeR’s (v3.32.1) calcNormFactors function (Robinson et al., 2010). For differential gene expression analysis, normalization and transformation were performed with Limma’s (v3.46.0) voom function (Ritchie et al., 2015, Law et al., 2014). Differential gene testing was performed with Limma’s lmFit, makeContrasts, contrasts.fit, and eBayes functions. For visualization purposes, this data was log2 CPM transformed using EdgeR’s cpm function and parameters: prior = 1, log = TRUE, normalized.lib.sizes = TRUE. Correspondence analyses were performed with MADE4 v1.64.0 (Culhane et al., 2005). For all heatmap visualisations and where required, sample standardization was performed by normalization to the mean expression of each gene. K-means clustering was performed with R’s (v4.0.2) base function kmeans with parameters: centers = 6, nstart = 25. K-means clustering was performed on the standardized log2CPM data (which was averaged between replicates prior to standardization). Hierarchical clustering was performed utilizing base R’s package stats (functions: dist and hclust), with the distance measure canberra and linkage method Ward.D. A set seed of 123 was used. Dendrogram visualization was performed with dendexted v1.15.1 (parameter: k = 3) (Galili, 2015); 3D visualizations were performed with plotly v4.9.4.1 (Sievert, 2020); heatmap visualizations were performed with ComplexHeatmap v2.6.2 (Gu et al., 2016); all other visualizations were performed with ggplot2 v3.3.5 (Villanueva and Chen, 2019) and where required ggrepel v0.9.1 (Slowikowski et al., 2018). Gene ontology and pathway analyses were performed with Metascape (http://metascape.org) (Zhou et al., 2019).

### Gene expression analysis of the human SARS-CoV2 genome

To quantify the amount of expression of hSARS-CoV2 across all samples, the raw counts data was utilized which included genes from both the human and hSARS-CoV2 genes. The raw counts data was processed and visualised by the same procedures as stated above (Gene expression analyses of the human genome). Specifically, it filtered, normalization factors were calculated, log2 CPM counts and CPM (parameter: log = FALSE) counts were generated, as well as standardized expressions. For visualization purposes, the expression of hSARS-CoV2 genes across the respective genome were ordered by the genomic feature’s starting base pair position.

### Custom genome for mapping

As the libraries were generated with p(A) enrichment, to avoid multimapping of other genes with ACE2, we generated a custom GENCODE’s GRCh38.p13 genomic reference files, where we removed the gene BMX. Additionally, we generated a custom hSARS-CoV2 (NC_045512v2) genomic reference files based on SwissProt precursor sequences (before cleavage) and Uniprot protein products (after cleavage) annotations. A custom genome combining these human and hSARS-CoV2 genomes was generated.

The protein products for annotation included: nsp1, nsp2, nsp3, nsp4, 3CL-PRO, nsp6, nsp7, nsp8, nsp9, nsp10, Pol, Hel, ExoN, nsp15, nsp16, Spike protein S1, Spike protein S2, ORF3a, E, M, ORF6. ORF7a, ORF7b, ORF8, N, and ORF10.

## Supplemental Figures

**Figure S1.**
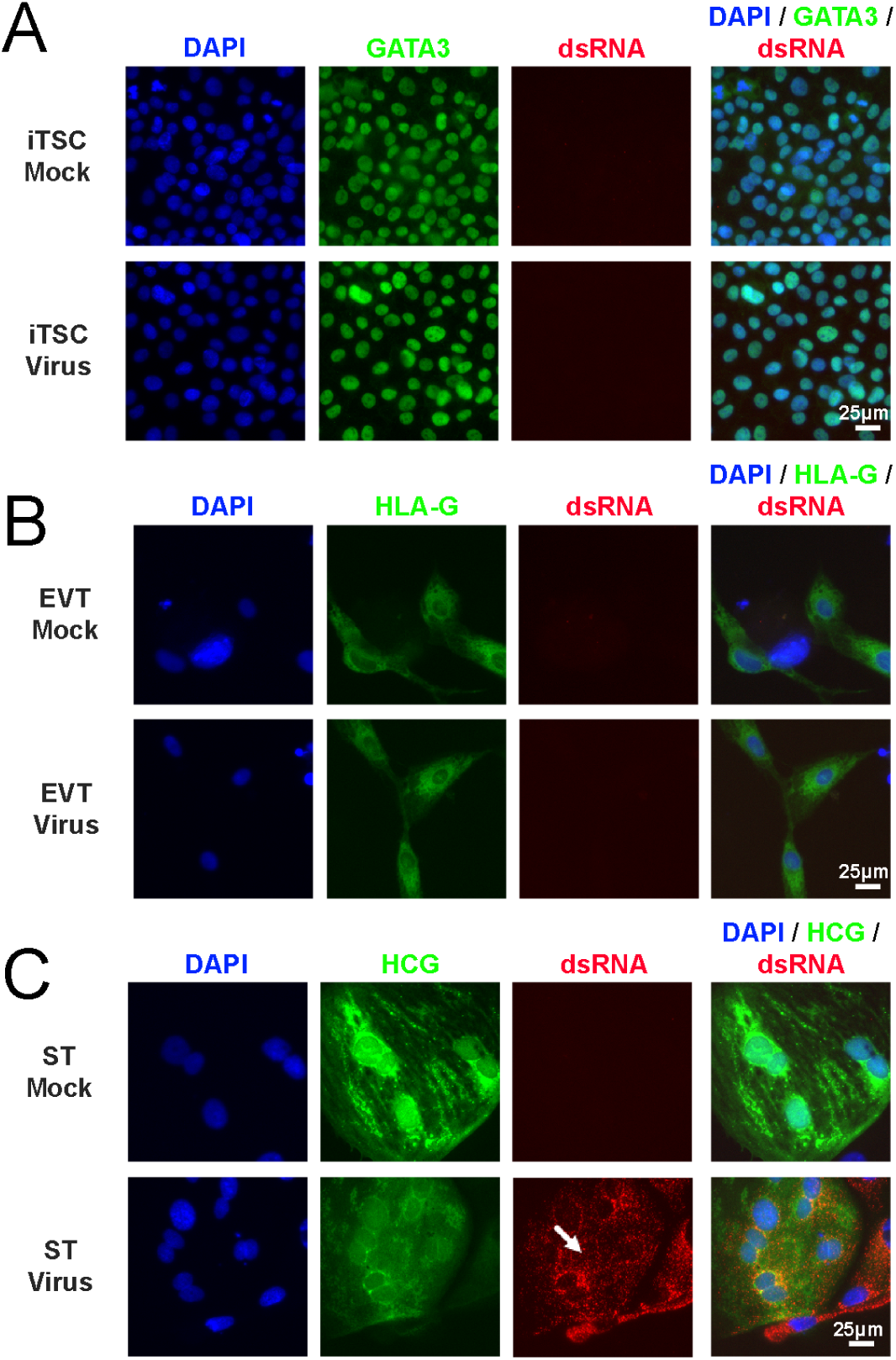
Immunofluorescence images of dsRNA (Red) along with GATA3 (A, TSCs; Green), HLA-G (B, EVTs; Green) or HCG (C, STs; Green) (32F cell lines). Cells were counterstained with DAPI.

**Figure S2.**
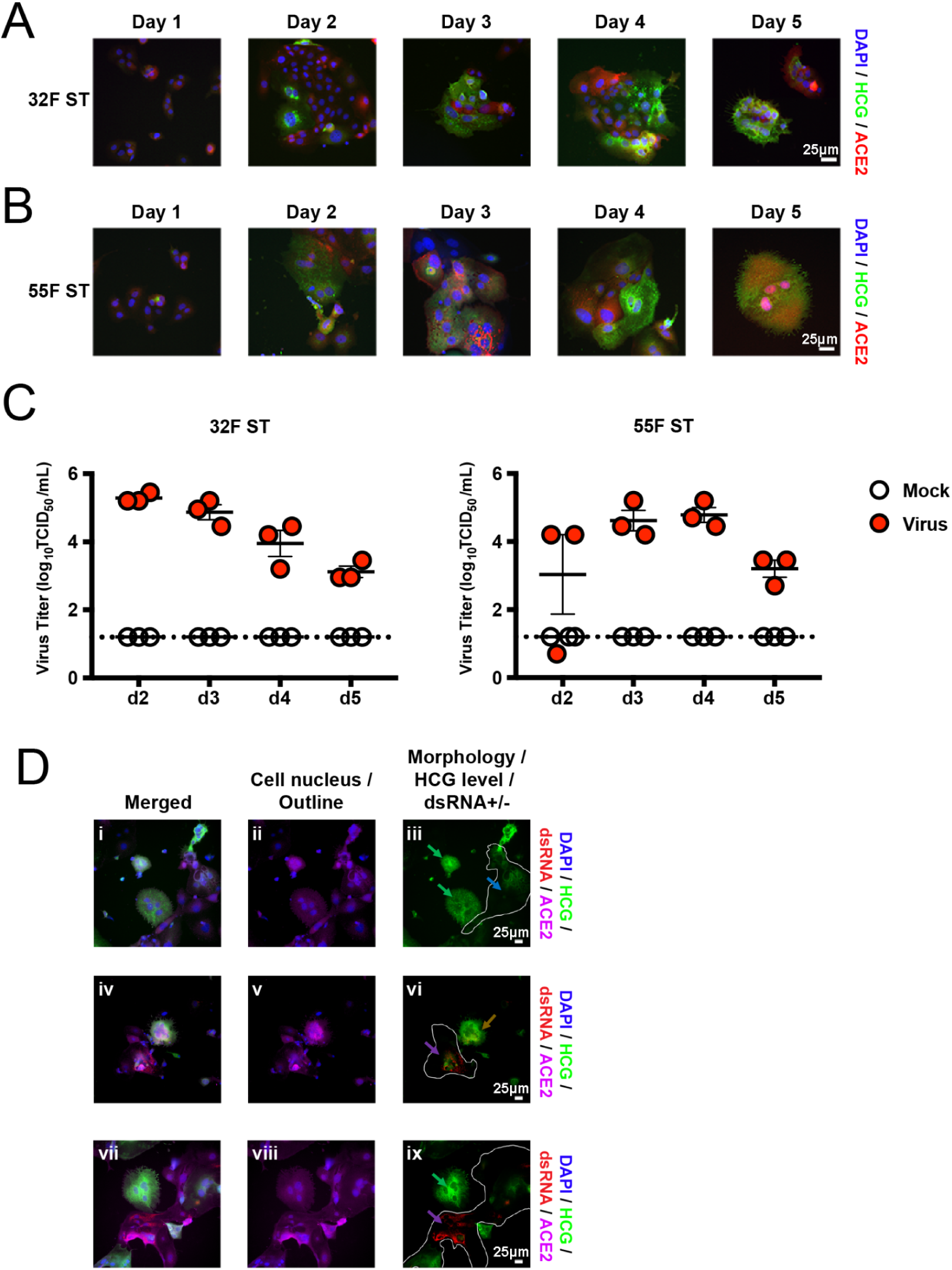
Immunofluorescence images of ACE2 (Red) along with , HLA-G (EVTs; Green) or HCG (STs; Green) on d1 to d6 (EVTs) and d1 to d5 (STs) in 32F cell lines (A) and 55F cell line (B). C) Virus titre expressed in log_10_TCID_50_/ml of 32F and 55F STs on d2 to d5. D) Scoring of morphology of STs. i, iv, vii) Merged images of DAPI (Blue), HCG (Green), dsRNA (Red), ACE2 (Magenta). ii, v, vii) Cell nucleus (DAPI) and outline of cells (ACE2) to mark multinucleated STs. iii) Merged HCG and dsRNA images for differentiated dsRNA- (green arrows) and undifferentiated dsRNA- (blue arrow, white outlined cell) cells. vi) Merged HCG and dsRNA images for differentiated dsRNA+ (brown arrow) and undifferentiated dsRNA+ (purple arrow, white outlined cell) cells. ix) Merged HCG and dsRNA images for differentiated dsRNA- (green arrow) and undifferentiated dsRNA+ (purple arrow, white outlined cell) cells. Scale bar=25µm. n=3 technical replicates of a representative experiment (C).

**Figure S3.**
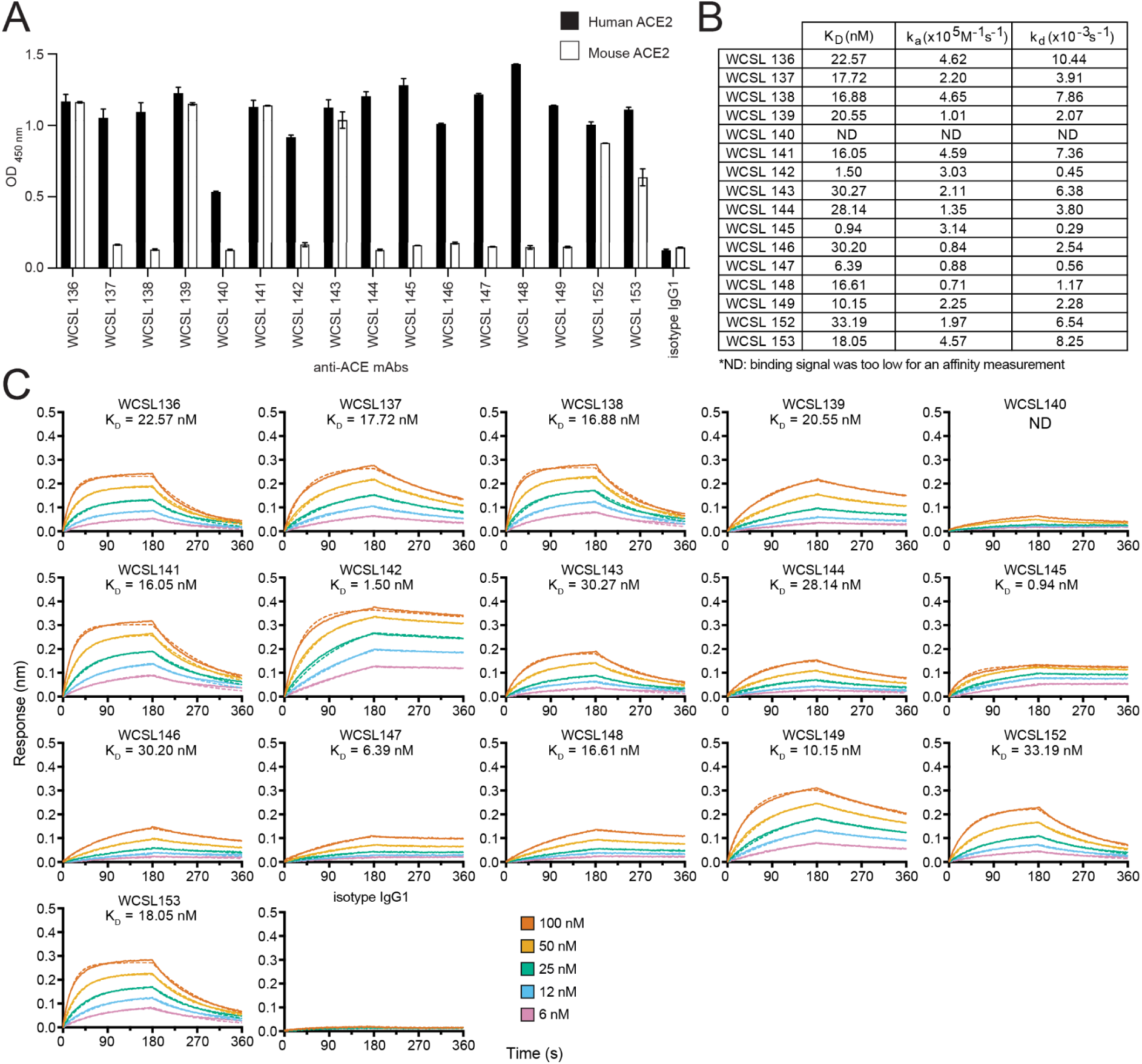
Recognition specificity of recombinant ACE2 and binding kinetics of anti-ACE2 monoclonal antibodies. (A) ELISA OD _450 nm_ signal of anti-ACE2 mAbs binding to human and mouse ACE2 (black and white bars respectively). Error bars represent mean ± standard deviation of technical duplicates. (B) Measured kinetic rate constants and affinity data for human ACE2 binding to immobilised anti-ACE2 mAbs. (C) Representative binding curves of five different human ACE2 concentrations from 6 - 100 nM binding to immobilised anti-ACE2 mAb. Responses measured from the experiment are represented by the solid line, and curves globally fit with a 1:1 binding model are represented by the dotted line. The isotype IgG1 control does not recognise ACE2.

**Figure S4.**
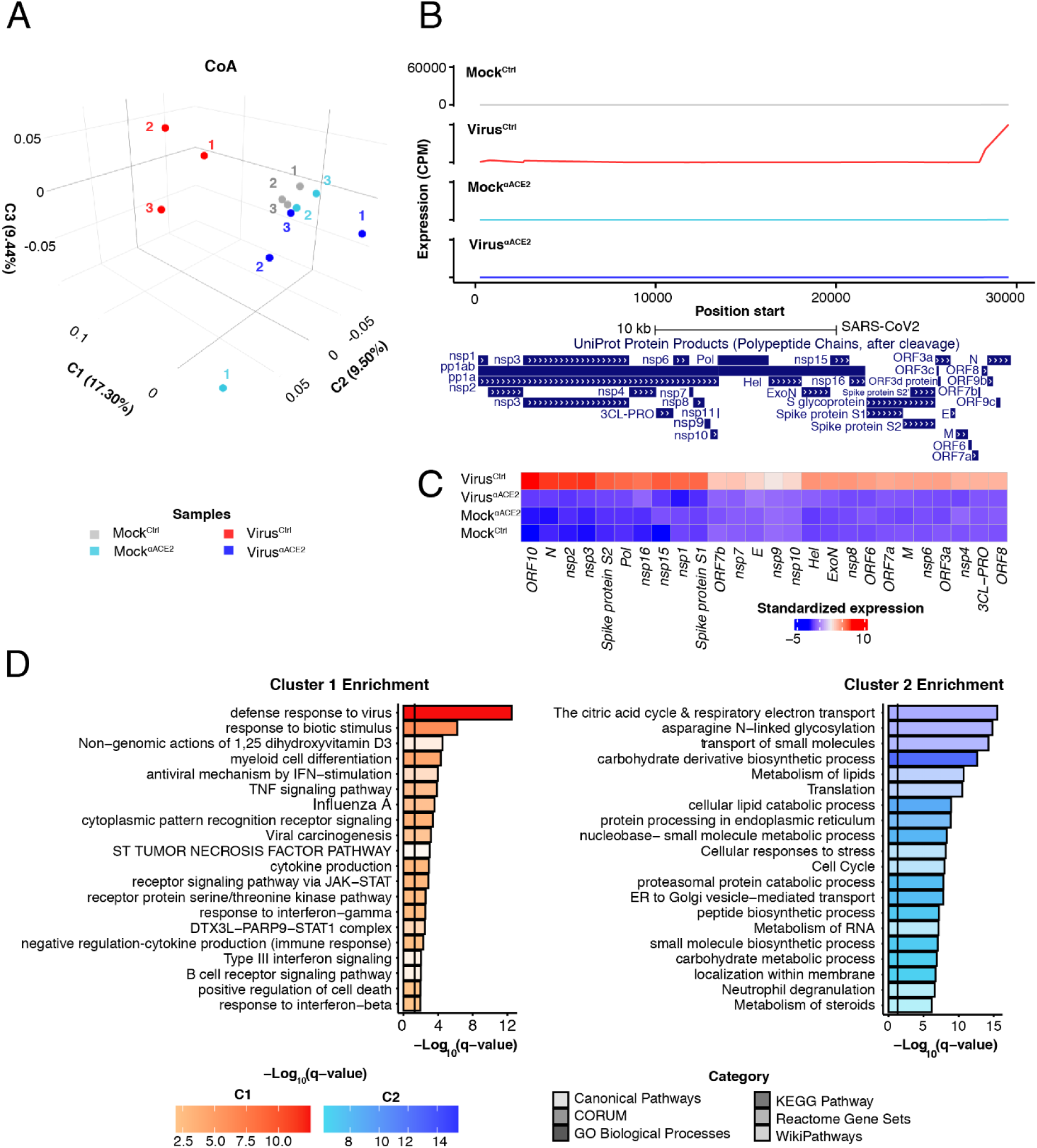
Transcriptome wide differences between STs under virus, mock and treated conditions, as well as hSARS-CoV-2 expression in the ST samples. A) Transcriptome-wide CoA of virus, mock and treated conditions of d3 infected STs. B) RNA expression levels of hSARS-CoV-2 genomic elements in d3 STs under virus, mock, and treated conditions. C) Hierarchically clustered heatmap of expression levels across the SARS-CoV2 genomic elements for d3 STs under virus, mock and treated conditions (dendrograms of clustering not shown). D) Functional enrichment analysis for genes upregulated (k-means cluster 1) and downregulated (k-means cluster 2) in STs infected with virus.

## Supplemental table legends

**Table S1. RNA-seq analysis outputs. DGE analysis output of virus vs mock samples related to** **Fig 3** **(sheet 1). K-means clustering analysis output of infected and mock samples, related to** **Fig 4** **(sheet 2).**

